# Communication subspaces align with training in ANNs

**DOI:** 10.1101/2024.11.11.623065

**Authors:** Peter G. Poggi, Stefan Mihalas, Dana Mastrovito

## Abstract

Communication subspaces have recently been identified as a promising mechanism for selectively routing information between brain areas. In this study, we explored whether communication sub-spaces develop with training in artificial neural networks (ANNs) and explored differences across connection types. Specifically, we analyzed the subspace angles between activations and weights in ResNet-50 before and after training. We found that activations were more aligned to the weight layers after training, although this effect decreased in deeper layers. We also analyzed the angles between pairs of weight layers. We found that for all branching, direct, and skip connections, weight layer pairs were more geometrically aligned in trained versus untrained models throughout the entire network. These findings indicate that such alignment is essential for the proper functioning of deep networks and highlights the potential to enhance training efficiency through pre-alignment. In biological data, our results motivate further exploration into whether learning induces similar subspace alignment.

## 1. Introduction

Given the brain’s vast complexity at many scales, understanding how information is communicated across networks, areas, neurons, and synapses has been quite challenging (Seguin et al., 2023). One of the main questions pertains to how populations of neurons selectively route information between brain regions that receive many inputs and have multiple outputs (Wang and Yang, 2018). Communication subspaces have emerged as one candidate mechanism for this selective routing. Communication subspaces can be conceptualized as low-dimensional projections of neuronal activity, capturing essential activity patterns of functional relevance. The principal directions of these lower-dimensional manifolds may dynamically rotate to achieve geometric alignment (Iyer et al., 2021), facilitating efficient information transfer between brain areas. Some evidence that the brain may utilize communication subspaces to selectively route information has been found in several recent studies; for instance, (Semedo et al., 2019) which first identified communication subspaces between macaque V1 and V2 in the context of a visual discrimination task and one study by Kaufman et al. (2014) which found that preparatory motor responses lie in a ‘null space’ while motor responses align with an ‘output-potent’ space.

As artificial neural networks often provide useful models for understanding principles of neural computation, we wondered whether evidence for communication subspaces could be found in them. Specifically, we were interested in whether geometrical alignment between layers increases as a function of training and how alignment differs in different connection types. To do this, we computed the angles between the singular vectors spanning the input and weight layers as well as those between pairs of weight layers in convolutional neural networks (CNNs). We focused our work on ResNet-50 for its architecture, consisting of multiple branching connections and skip connections (see He et al. (2016) for more about ResNet architectures).

## 2. Methods

### 2.1. Computing the Angles between Subspaces

#### Computing alignment between convolutional weight layers involves several steps (Figure 1)

1) First, we reorganize the convolutional weights into 2D connection matrices that can be applied to inputs via multiplication rather than convolution. This was achieved by flattening and reorganizing the input features and kernels such that each row in the weight matrix corresponds to the flattened kernels of a single output channel and each column in the input matrix represents the patches of input features that align with the convolution operation for an output channel as shown in A3 (Liu et al., 2019). The resulting matrix has output channels along the rows and combined input channels and kernel dimensions across the columns.

**Figure 1.**
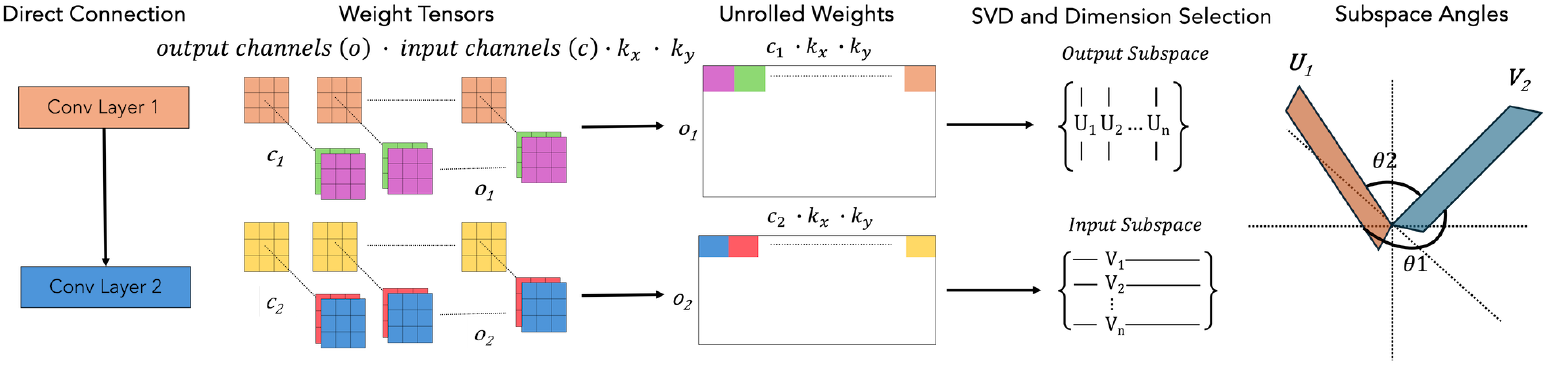
Schematic representation of the steps to compute subspace angles between weight layers in a direct connection.
2) Second, we decompose the resultant 2D connection matrices into their constituent singular vectors using SVD. When decomposing the weight matrix, the left singular vectors span the output channel dimension resulting in an output space while the right singular vectors span the input channel and kernel dimensions yielding an input space. Although our analysis took place in feature space due to prohibitively large neuron connectivity (Toeplitz) matrices in modern CNNs, ResNet’s 1×1 bottleneck kernels ensured that the input and output spaces had the same dimensionality, making our method a good approximation of the full Toeplitz matrix.
3) Finally, we compute the angles between the singular vectors. To measure alignment, we used the principal or canonical angles between subspaces which represent the canonical correlations of matrix pairs between the column space of two flats. Given two subspaces *X* ∈ *R*^*n×p*^ and *Y* ∈ *R*^*m×q*^, there will be *z* principal angles, where *z* = min(*p, q*) (see Knyazev and Argentati, 2002 for a mathematical overview of principle angles). We chose our subspace dimensions based on the maximum amount of explained variance we could attain while excluding extremely small angles within the null space of the matrices. This resulted in subspaces retaining around 50% of the dimensions and *>*70% of the explained variance of the original input and output spaces. We then took the mean of the principal angles 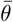, which gave us a single measure over multidimensional subspaces.

### 2.2. Training

Pre-training weights were randomly initialized from a Kaiming normal distribution. Trained weights for ResNet-50 were obtained from torchvision.models. The model used was trained to 76.13% accuracy on the ImageNet1K dataset.

### 2.3. Connection Types

We analyzed five different connection types in ResNet-50: direct connections, branching connections, short and long skip connections, and indirect connections (see A4 for diagrams). The simplest is a direct connection where the output of a convolutional layer is directly fed to the next convolutional layer. The second type of connection is the branching connection where a certain layer sends the same output to two different convolutional layers. There are also skip connections implemented by element-wise addition whose output and input weight layers can be compared. There are two possible arrangements for this, one which compares the output of the last convolutional layer with the input of the first convolutional layer of the next block (short skip) and another which compares the first layers of each block to each other (long skip). Finally, as a control, we compared the alignment between layers and inputs that were indirectly connected (via connections to intermediate convolutional layers) and whose activity did not belong to the same space. Thus, we did not expect alignment to develop during training.

### 2.4. Measuring Activity Alignment

Activity, in the form of activation feature maps, represents inputs to convolutional layers. As such, we used their left singular vectors to characterize their activity. In all cases, we compared the left singular vectors of the activations to the right singular vectors of the weight layers (the input space). Our inputs consisted of 100 random images from the ImageNet 2012 validation set. We calculated the alignment between activations and weights individually for all connection types.

### 2.5. Measuring Layer Alignment

To measure layer alignment, we compared the left singular vectors of a given layer (output space) to the right singular vectors (input space) of the subsequent layer in all connection types except for branching. For branching we compared the two input spaces of the branches to each other which told us how aligned the branches were to each other.

## 3. Results

We performed two distinct analyses. The first tested the alignment between activations and weight layers while the second tested the alignment between the weight layers themselves without any inputs. All code can be found at NeurReps 2024 Communication Subspaces.

### 3.1. Activation-to-weight alignment

We found that activations were more aligned in the trained model across all layers for branching, short skip, and long skip connections (Fig.2b-d, Mann-Whitney U test; p *<*.001. See B5 for exact p-values). Direct connection layers were significantly more aligned until the last three blocks Fig.2a. This likely reflects an increased selectivity of information flow in later layers as the network comes closer to classifying the image. The skip connections were less aligned than direct and branching connections due to the rotation of the space introduced by intermediate layers which are added to the output. Nevertheless, the trained layers remained significantly more aligned than the untrained layers, demonstrating the effectiveness of the residual connections in maintaining alignment across multiple layers. As expected, none of the indirect connections were significantly different between untrained and trained models Fig.2e.

### 3.2. Weight-to-weight alignment

The alignment of weight layers closely resembled that of the activation-weight layer alignment (Figure 2-bottom). Trained weight layers were significantly more aligned than untrained layers across connection types (Fig.2f-i, Mann-Whitney U test; p *<*.05) except for indirect connections (control). As with activation-weight layer alignment, direct connections and branching connections were more aligned than skip connections. However, unlike the alignment of the activation-weight layers, we did not find a decrease in alignment with network depth. This suggests that weight alignment may be necessary throughout the network to maintain functional connections.

**Figure 2.**
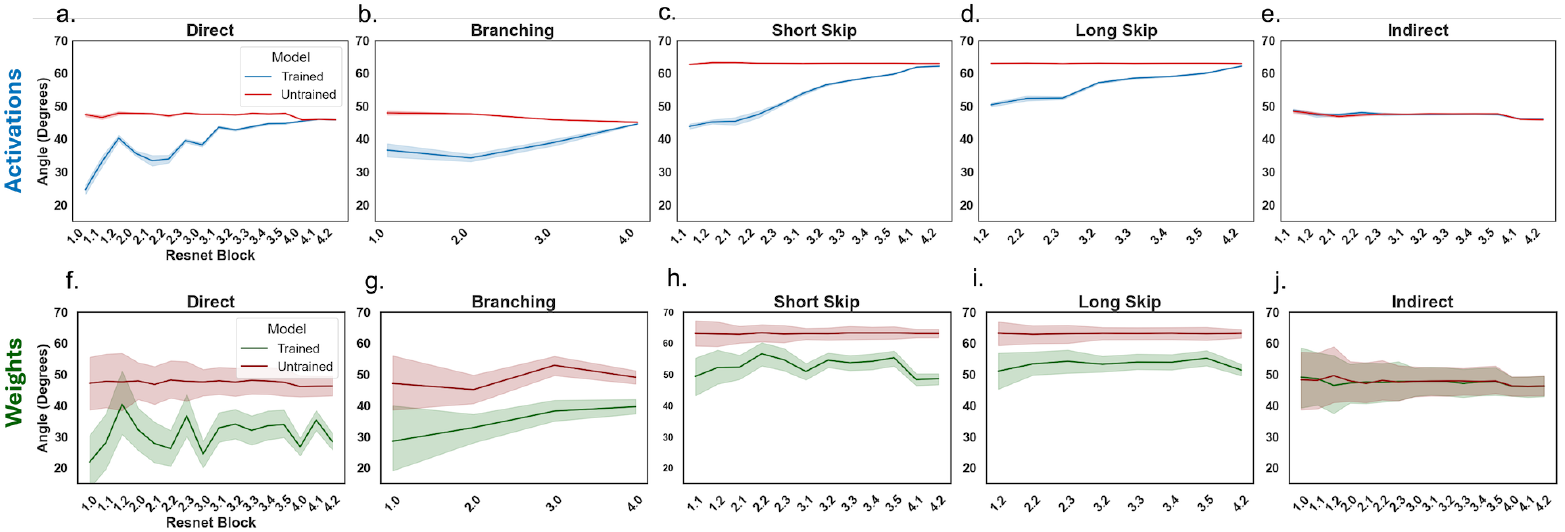
Top: mean alignment between activation-to-weight connections. Bottom: mean alignment for weight-to-weight connections across layers.

## 4. Conclusion

Our study found that communication subspaces develop and become aligned in ANNs over training, with connection-specific differences. This finding supports the hypothesis that training may improve inter-areal communication through subspace alignment, a theory we are currently investigating using biological data in a visual change-detection and familiarity task. In contrast to Iyer et al. (2021), who found that subspaces in mouse visual cortex are relatively unaligned until stimuli are presented whereby they briefly align; CNNs maintain alignment throughout as their only function is image classification. Our work provides a framework for investigating selective routing between different types of connections in the brain, especially in branching structures. Additionally, we see potential applications in machine learning, where pre-alignment may be employed to improve training efficiency in ANNs. Together, this study lays the groundwork for further research into inter-areal communication and selective information routing via subspace alignment, offering further avenues for both biological exploration and neural network training.

## 5. Acknowledgement

We would like to thank Paul G. Allen for his funding and support.

## Appendix A.

**Figure 3.**
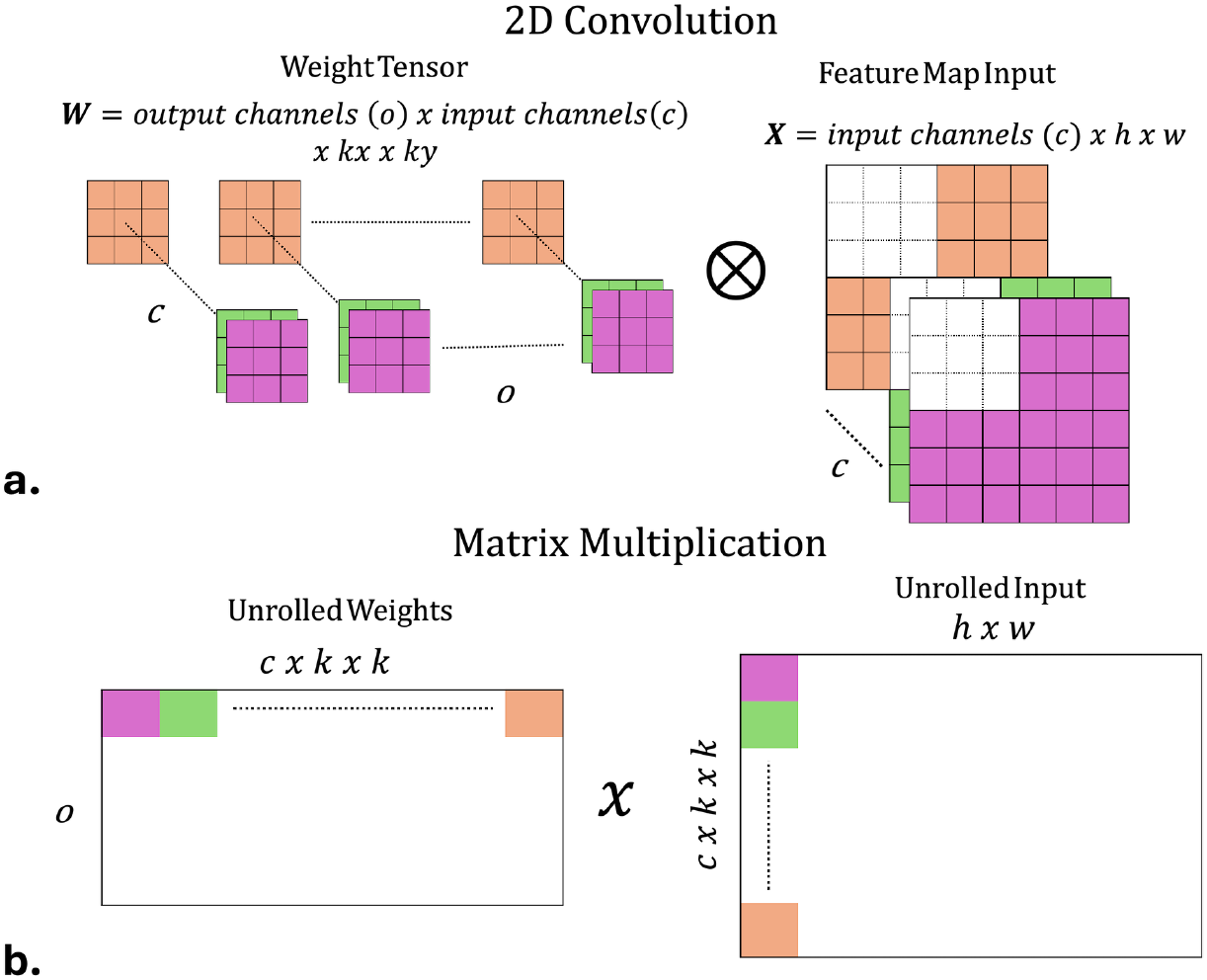
Panel a. visually depicts normal 2D convolution. Panel b. depicts the matrix multiplication equivalent as in Liu et al., 2019.

**Figure 4.**
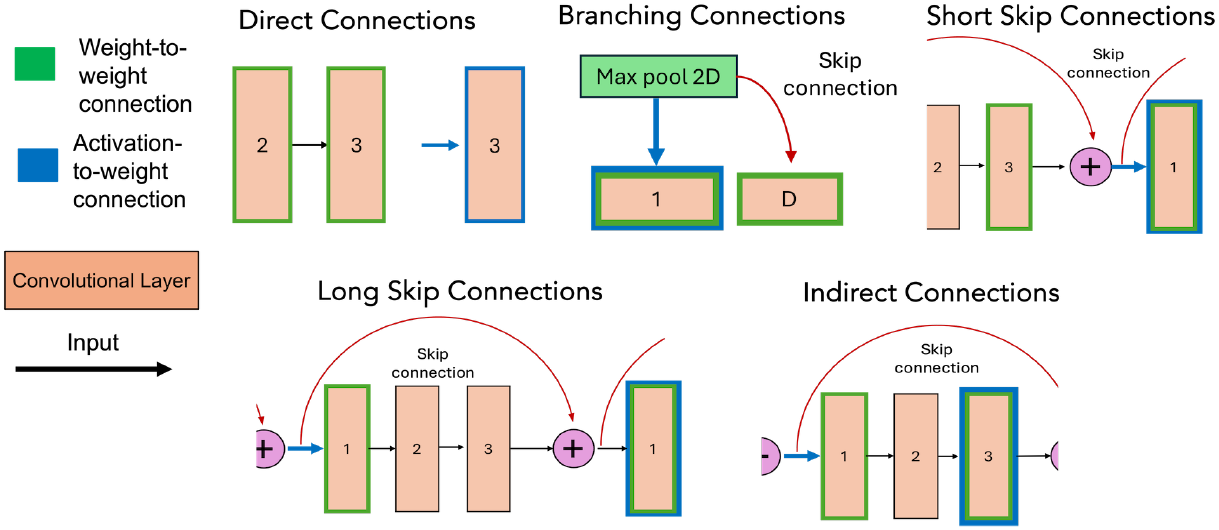
Diagramatic representation of various connection types for weight-to-weight and activation-to-weight connections.

## Appendix B.

**Figure 5.**
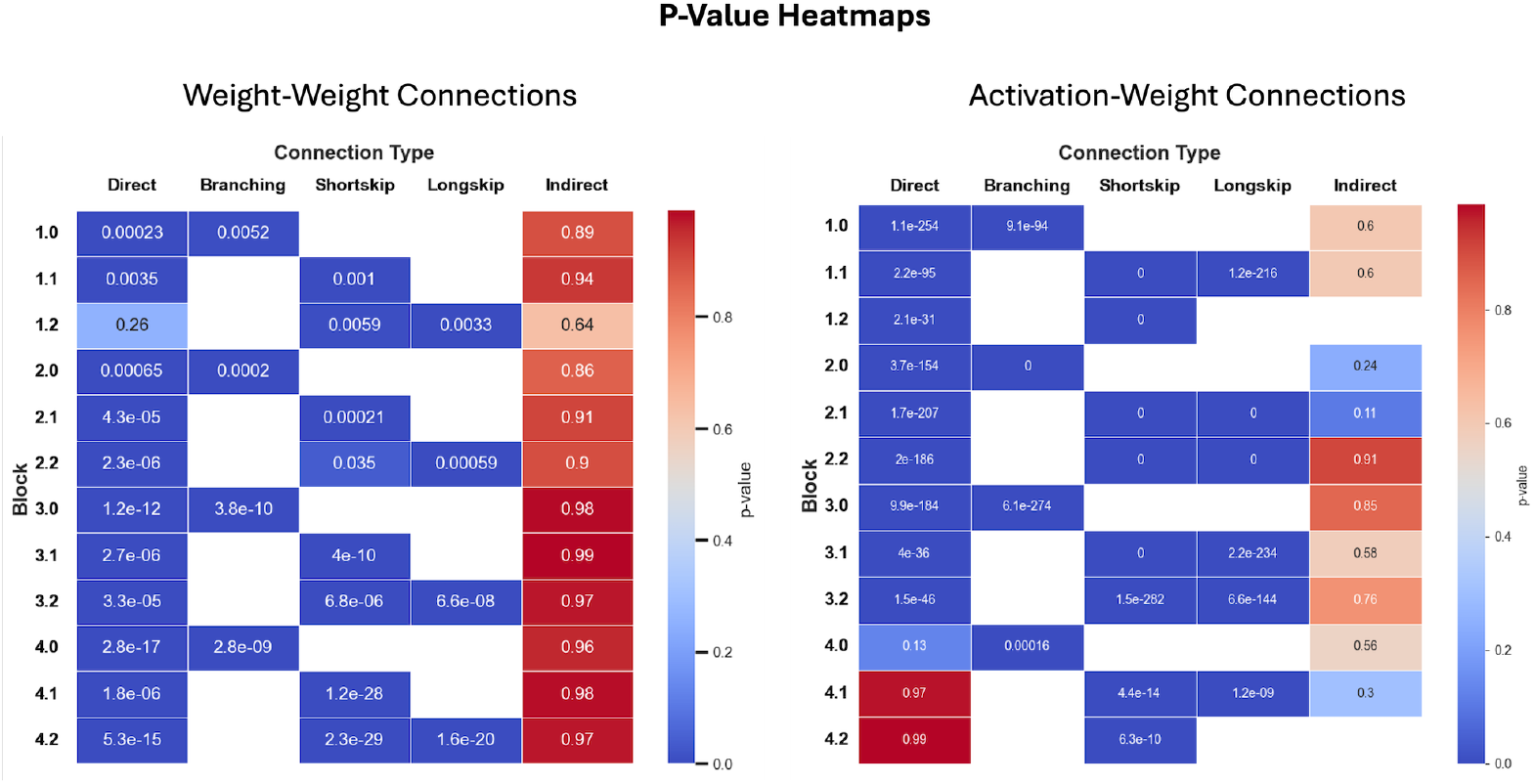
Heat maps of the exact p-values for each connection type and layer for both analyses. All dark blue squares are statistically significant (Mann-Whitney U test; p *<*.05). Blank spots indicate that either there is no connection of that type in the given block, or the output and input spaces were not of the same row dimensions and could not be compared (e.g. branching only occurs in layers 1.0, 2.0, 3.0, and 4.0 so the branching results are only for these blocks).

